# Divergent selection predating the Last Glacial Maximum mainly acted on macro-phenotypes in Norway spruce

**DOI:** 10.1101/2022.01.15.476468

**Authors:** M. Tiret, L. Olsson, T. Grahn, B. Karlsson, P. Milesi, M. Lascoux, S.-O. Lundqvist, M.R. García-Gil

**Author notes:** corresponding author: Mathieu Tiret, **Email:**.

## Abstract

The current distribution and population structure of many species were, to a large extent, shaped by cycles of isolation in glacial refugia and subsequent population expansions. Isolation in, and post-glacial expansion through heterogeneous environments led to either neutral or adaptive divergence. Norway spruce is no exception, and its current distribution is the consequence of a constant interplay between evolutionary and demographic processes. We investigated population differentiation and adaptation of Norway spruce for juvenile growth, diameter of the stem, wood density and tracheid traits at breast height. Data from 4,461 phenotyped and genotyped Norway spruce from 396 half-sib families in two progeny tests were used to test for divergent selection in the framework of Q_ST_ vs F_ST_. We show that the macroscopic resultant trait (stem diameter), unlike its microscopic components (tracheid dimensions) and juvenile growth, was under divergent selection that predated the Last Glacial Maximum. Altogether, the current variation in these phenotypic traits in Norway spruce is better explained by local adaptation to ancestral environments than to current ones, where populations were partly pre-adapted, mainly through growth-related traits.

## Introduction

During the Quaternary glacial cycles (up to 2.5 million years ago), many European animal and plant species were repeatedly confined to southern refugia. Those were often located in the Iberian, Italian and Balkan peninsulas, but surviving populations could also be distributed on less discrete spots and, instead, scattered along the southern edge of the ice sheet (Bennett, 1997; Binney et al., 2009; Kremer et al., 2010). Improvement of climatic conditions during post-glacial periods led to demographic expansions accompanied by population admixture (e.g., Petit et al., 2003), new selective pressures (e.g., Suarez & Tsuitsui, 2008), and inter and/or intraspecific competition (e.g., Savolainen et al., 2007). Although the present population structure is the result of successive cycles of glaciation events that led to a series of demographic contractions and expansions, the current distribution of many species has, at least in part, been shaped by the different ecological processes and selective pressures that acted during the period after the Last Glacial Maximum (e.g., Petit et al., 2002; Palmé et al., 2003).

New abiotic and biotic pressures encountered during the glacial cycles presumably led to new adaptations. In many tree species, current phenotypes partly reflect past selection (Von Gadow et al., 2021), and selection is known to play an important role in shaping species distribution (Case et al., 2005; Savolainen et al., 2007; Wisz et al., 2013; Li et al., 2022). Among others, wood is one of the main tissues undergoing selection, since radial growth of the woody stem provides the mechanical support for woody plants to grow tall and gain a competitive advantage (Givnish, 1982; Iwasa, 1985; Falster & Westoby, 2003; Falster & Westoby, 2005). However, adaptation for upright growth is often associated with modified wood properties and evolutionary trade-offs (Poorter et al., 2006; Chave et al., 2009). Wood density is a good example of such a growth-survival trade-off (Kitajima 1994; Poorter et al., 2010), where low wood density, resulting from wider lumina and, in hardwood species, wider vessels (e.g., Enquist et al., 1999; Preston et al., 2006; Coomes et al., 2008), is associated with fast growth. Conversely, high wood density provides higher biomechanical safety (e.g., Niklas, 1993; Van Gelder et al., 2006) and resistance against pathogens (Augspurger & Kelly, 1984), and is thus associated with higher survival rates. The growth-survival trade-off is a common ecological trade-off observed in both tropical (e.g., Faster & Westoby, 2005; Poorter, 2008) and boreal species (e.g., Rossi et al., 2015; Pretzsch et al., 2018).

Norway spruce (*Picea abies* (L.) H. Karst.) is a dominant boreal conifer species that thrives nowadays in a vast northern domain spanning from Norway to central Russia, as well as in smaller European domains, such as the Alps and the Carpathians (Vendramin et al., 2000; Tsuda et al., 2016; Li et al., 2022; Nota et al., 2022). Repeated demographic movements associated to climatic cycles and ensuing secondary contacts led to the division of Norway spruce into seven genetic clusters (Chen et al., 2019): Northern Fennoscandia (NFE), Central and South Sweden (CSE), Russia-Baltics (RBA), Northern Poland (NPL), Central Europe (CEU), Alpine (ALP), Carpathian (ROM) domains. In southern Sweden, recent hybrids between CSE and ALP clusters can be encountered (CSE-ALP). The demographic history leading to this population structure went along a history of successive adaptations to biotic and abiotic factors. Following the hypothesis that most adaptation is reflected by cluster specific allele frequencies (Kremer et al., 2012), we investigated past adaptive signature through the population structure of Norway spruce. Using the framework of Q_ST_ vs F_ST_, we assessed whether a phenotype diverges more between the genetic clusters than expected under random genetic drift. The timing of the divergence thereby inferred was given by that of the population structure that long predated the Last Glacial Maximum (10 to 15 million years ago; Chen et al., 2019; Li et al., 2022). By doing so, we were able to infer the proportion of the current phenotypic variation that is due to adaptations predating the Last Glacial Maximum. For that, in order to account for possible evolutionary trade-offs between wood-related traits, we analyzed several phenotypes: stem diameter and ring widths, wood density and tracheid traits defining it. Since the nature of selection between trees is expected to change over ontogenetic stages (Miriti, 2006), we studied the development of these phenotypes with cambial age/ring number from increment cores sampled from two progeny trials of the Swedish breeding program. We have conducted the analyses on a large dataset of 4,461 twenty-year-old trees from 396 half-sib families, 118,145 SNPs from exome sequencing. We show that the current phenotypic variation is partly explained by divergent selection that predated the Last Glacial Maximum, which mainly acted on macro-phenotypes (stem diameter), rather than micro-phenotypes (tracheid dimensions).

## Materials and Methods

### Plant material

The study is based on previously presented data (Chen et al., 2014; Chen et al., 2016; Lundqvist et al., 2018), which are here analyzed with a special attention to population structure and selection. The data originates from two progeny trials of Norway spruce at two different sites: FF1146 (Höreda, Sweden; 57°60’N 15°04’E), and FF1147 (Erikstorp, Sweden; 56°80’N 13°90’E). Both trials were established by the Forestry Research Institute of Sweden (Skogforsk) in the spring of 1990 from open-pollinated seeds harvested on selected trees from 112 stands in southern Sweden. FF1146 comprised 1,373 half-sib families and FF1147, 1,375 half-sib families. In both trials, a randomized incomplete block design was implemented (20 blocks in FF1146, 23 in FF1147), with single-tree plots and a homogeneous spacing of 1.4 m x 1.4 m. Dead trees and trees of low breeding interest were removed in 2008. After filtering out (i) half-sib families absent in the established Skogforsk clonal archives, and (ii) stands with less than three half-sib families, the study included a total of 518 half-sib families with 5,644 progeny (2,983 from FF1146, and 2,661 from FF1147).

### Population structure

Population structure was previously assessed at a fine scale using 399,801 unlinked putatively non-coding SNPs on 1,672 individuals (see details in Chen et al., 2019), categorizing families into the following genetic clusters (ordered by decreasing latitude): Central and South Sweden (CSE), Russia-Baltics (RBA), Northern Poland (NPL), Central Europe (CEU), Alps (ALP), Carpathians (ROM), and hybrids between CSE and ALP (CSE-ALP) (for tree geographical origins, see Fig. 1 in Milesi et al., 2019). The families examined in this study followed the same genetic clustering (Fig. 1A, 1B). Individuals from the ROM genetic cluster were removed, as our sample included only three such families, whereas all other clusters included at least ten families (Table S1).

**Figure 1.**
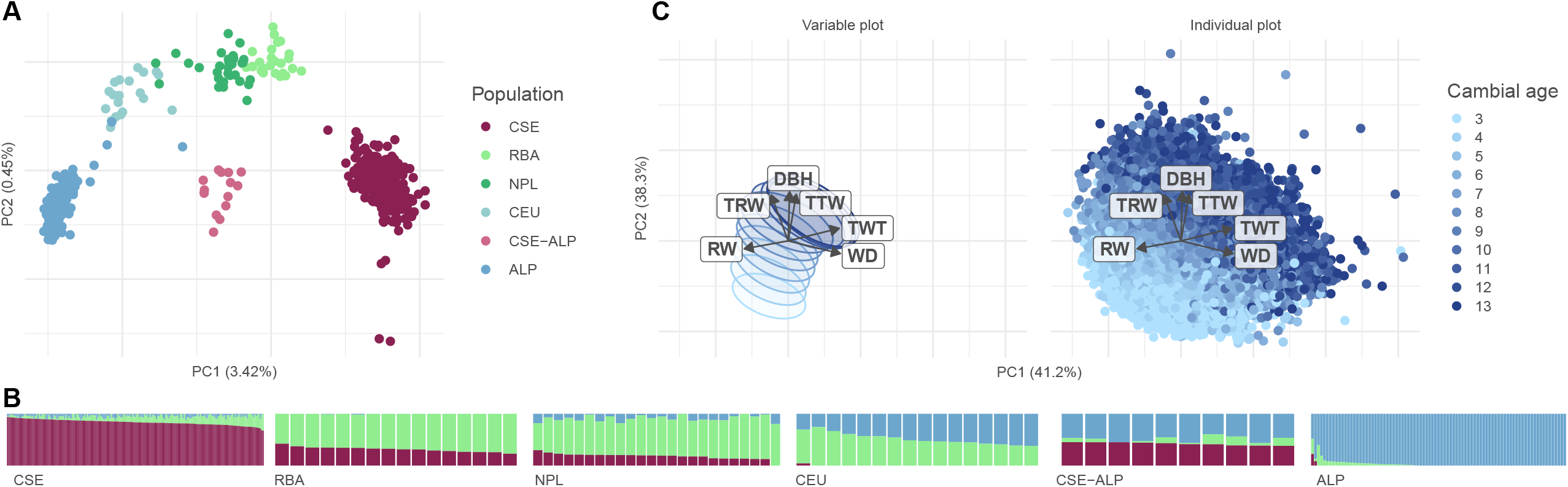
Overall description of phenotypes and genotypes of the sample of Norway spruce. **(A)** Principal Component Analysis of the genotypes of the half-sib families. Families are colored according to their genetic clusters (ordered by decreasing latitude in the legend): Central and South Sweden (CSE, dark red), Russia-Baltics (RBA, light green), Northern Poland (NPL, green), Central Europe (CEU, light blue), Alps (ALP, blue), and hybrids between CSE and ALP (CSE-ALP, pink). PC1 and PC2 explained 3.42% and 0.45% of the variance, respectively. **(B)** Admixture plot of the half-sib families, with K = 3. Individuals are grouped by population, and sorted according to their coancestry coefficient. **(C)** Principal Component Analysis of the phenotypes: Diameter at Breast Height (DBH), Ring Width (RW), Wood Density (WD), Tracheid Radial Width (TRW), Tracheid Tangential Width (TTW), and Tracheid Wall Thickness (TWT). Left panel only shows variable axes and 60% confidence ellipses. On the right panel, each data (dot) is colored according to its cambial age (3^rd^ year in light blue up until 13^th^ year in dark blue). PC1 and PC2 explained 41.2% and 38.3% of the variance, respectively.

### Phenotype measurements

Increment cores of 12 mm diameter, extracted from the northern side of the stem, were collected in 2010 and 2011 at breast height (BH, 1.3 m above ground) from six trees per half-sib family per trial. The usual cutting age is 60-70 years or more, while the cores were extracted from 20- or 21-year-old trees: only juvenile stages of wood characteristic were then captured by the cores. The increment cores were analyzed by SilviScan (Evans, 1994; 2006) at Innventia, now RISE (Stockholm, Sweden), assessing high-resolution pith-to-bark radial variation in growth, tracheid and wood properties. In this study, we focused on the development with cambial age (ring number) of ring width (RW, in mm), and averages per ring of wood density (WD, in kg/m^3^), tracheid radial width (TRW, in μm), tracheid tangential width (TTW, in μm), tracheid wall thickness (TWT, in μm), diameter at breast height (DBH, in mm) calculated from ring widths, and the age when the trees reached breast height (ABH). Although ABH is related to upright growth, it can be estimated accurately with high quality increment cores from old trees. The measurements, identification of annual rings, and the importance of ABH are discussed in Lundqvist et al. (2018).

The two innermost rings were removed as the extremely curved structure of the xylem does not allow the same high data quality as for rings further out. In addition, only data up to the 13^th^ ring was kept for analyses, since only fast-growing offspring with low ABH are represented at the highest cambial ages. To avoid the effect of the thinning performed, we removed all annual rings formed after 2008. The study then covered six traits (RW, WD, TRW, TTW, TWT, DBH) for 11 rings per offspring in most of the cases, and their ABH.

### Exome capture and SNP filtering

The exome capture (target re-sequencing) of the mother of each half-sib family is available on BioProject PRJNA511374 (Chen et al., 2019) and BioProject PRJNA731384 (Chen et al., 2021). Total genomic data was extracted from the buds of the 518 maternal trees (or needles when buds were not available) using the Qiagen Plant DNA extraction kit (Qiagen, Germany). DNA concentration was assessed using the Qubit® ds DNA Broad Range Assay Kit (Oregon, USA). The extracted DNA was sent to RAPiD Genomics (Gainesville, USA) for library preparation and exome sequencing. Sequence capture, with an average depth of 15X, was performed using the Illumina HiSeq 2500 (San Diego, USA), with 40,018 diploid probes (120bp-length) designed to capture 26,219 genes of *P. abies* (Vidalis et al., 2018). After mapping the raw reads against the *P. abies* reference genome v1.0 (Nystedt et al., 2013) using BWA-MEM with default settings (Li & Durbin, 2010), PCR duplicates were sorted and marked using SAMTOOLS v1.2 (Li et al., 2009) and Picard v1.141 (http://broadinstitute.github.io/picard). INDELs were realigned using GATK v3.5.0 (McKenna et al., 2010). Variant calling was performed using GATK HaplotypeCaller v3.6, non-biallelic SNPs were filtered out using GATK SelectVariants v3.6 (option --restrict-alleles-to BIALLELIC), and quality filtering was carried out by the Variant Quality Score Recalibration following the procedure of Van der Auwera et al. (2013).

As an alternative way to account for Minor Allele Frequency (MAF), we filtered out SNPs with less than five individuals per genotype to avoid a leverage effect. We performed a loose filtering for variant call rate (or missingness) with PLINK v1.9 (Chang et al., 2015), by removing individuals with more than 70% SNP-missingness, resulting in 472 half-sib families, and by removing SNPs with more than 70% individual-missingness, resulting in 118,145 SNPs (each mother is then genotyped for at least 35,443 SNPs). 70,277 SNPs were successfully mapped on the consensus map of Bernhardsson et al. (2019).

### Statistical model

The traits (DBH, WD, RW, TRW, TTW, and TWT) were normalized by their respective overall means (across all trees and rings 3 to 13) to allow comparison between traits. Then, they were analyzed at each cambial age separately, resulting in six traits over 11 rings, a total of 66 traits in addition to ABH.

To study the family effect underlying the traits, we used the following mixed model:

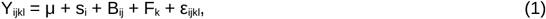

where ‘Y_ijkl_’ is the trait of the l-th offspring of the k-th family in the j-th block at the i-th site, ‘μ’ is the grand mean, ‘s_i_’ is the fixed effect of the i-th site, ‘B_ij_’ and ‘F_k_’ are uncorrelated random effects of the j-th block nested in i-th site (with variance σ_B_^2^) and the k-th half-sib family (with variance σ_F_^2^) respectively, and ‘ε_ijkl_’ is the residual fitted to a Gaussian distribution of variance σ^2^. Heritability was estimated as 4 x σ_F_^2^ / (σ_F_^2^ + σ_B_^2^ + σ^2^) as an intraclass correlation estimation of half-sib families (p.166, Falconer, 1989).

To investigate the effect of genetic clusters on traits jointly with the effect of ABH, we used the following linear mixed model:

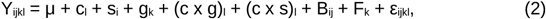

where ‘c_l_’ is the fixed effect of the l-th offspring’s ABH, ‘(c x g)_l_’ is the fixed interaction effect between ABH and the genetic cluster of the l-th offspring, ‘(c x s)_l_’ is the fixed interaction effect between ABH and the site of the l-th offspring, and the other terms are as defined above in models (1). Model (2) was applied on the 66 traits, as previously mentioned. Since the interaction between ABH and the site effect was always significant, we split the dataset per site, as recommended by Crawley (2007, pp. 323–386). The results are focusing on the Höreda trial, but the conclusions were the same in the Erikstorp trial.

Finally, to disentangle the effect of divergent selection and neutral divergence, we estimated Q_ST_ as defined by Spitze (1993) with the following mixed model:

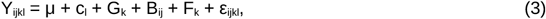

where ‘G_k_’ is the random effect of the k-th family’s genetic cluster (with variance σ_A_^2^), and the other terms are as defined above in model (1). Q_ST_ was estimated as σ_A_^2^/(σ_A_^2^ + 2σ_F_^2^), and averaged across rings. Following the recommendation of Whitlock (2008), we compared Q_ST_ to the positive part of the distribution of F_ST_ (with VCFTools v0.1.16, option --weir-fst-pop; Danecek et al., 2011) computed on SNPs in chromosomic regions identified as non-coding by Nystedt et al. (2013). This set was not filtered for MAF or missingness, amounting then to 789,786 SNPs. If Q_ST_ of a trait is significantly larger than F_ST_, the trait can be considered to have diversified more than expected under neutrality in a heterogeneous environment. In contrast, if Q_ST_ cannot be distinguished from F_ST_ values, we would have little evidence that selection is responsible for phenotypic divergence.

All computations were done using R v4.0.4 (R core team, 2021), and all models were fitted with the R package *lme4* (function *lmer*; Bates et al., 2015). Statistical significance of the fixed effects was assessed using a type II Wald chi-square test (function *Anova* of the R package *car*), and significance of a slope was assessed using a Student *t-*test (function *t*.*test* of the R package *stats*). To account for the unbalance caused by consecutive filtering, we only kept families with at least four offspring per site. Applying all the previous filters, we ended up with 396 half-sib families and 4,461 offspring per trait. Multiple testing across the 66 traits were accounted for by considering a Bonferroni adjusted cut-off of p = 7.6 × 10^−4^.

## Results

### The genetic cluster effect was significant at all ages

At all cambial ages, phenotypic variation of all six phenotypes (DBH, WD, RW, TRW, TTW, and TWT) was associated with the geographical origin of the genetic cluster, as the genetic cluster effect was significant for all rings (χ^2^ > 11.6, degree of freedom or df = 5, p = 0.041), except for the 3^rd^ ring TWT (χ^2^ = 10.9, df = 5, p = 0.053). Every trait of the southern populations was consistent with a higher investment in growth than in wood density (higher DBH, RW, TRW, and TTW; lower WD, and TWT).

Phenotypic variation exhibited a strong temporal pattern, an ontogenetic shift (Fig. 1C). For all rings, phenotypic variation also depended on the geographical origin of the trees. The ontogeny was quantitatively measurable, as it followed the second axis of the PCA on the six phenotypes (41.2% by PC1, 38.3% by PC2). As expected, the ontogeny was mainly driven by that of DBH, which alone explained 34% of PC2, and on average increased fourfold over the cambial ages investigated, while other phenotypes varied at most by 54% (for further details, see Lundqvist et al., 2018). As cambial age increased, ontogenetic shift was instead driven by other phenotypes, as the coefficient of variation of DBH was the only one decreasing over time (Student’s t = −6.13, p < 0.001; Fig. S1). Overall, the ontogenetic shift resulted from a complex interplay of multiple phenotypes, all with heritabilities higher than 30% (Table 1), in accordance with previously reported values (Hannrup et al., 2004; Chen et al., 2014, 2021).

**Table 1.** Genetic variance (σ_F_^2^), environmental variance (σ_B_^2^ + σ^2^), and heritability (h^2^) were estimated with model (1). Traits: Calendar age at Breast Height (ABH), Diameter at Breast Height (DBH), Ring Width (RW), Wood Density (WD), Tracheid Radial Width (TRW), Tracheid Tangential Width (TTW), and Tracheid Wall Thickness (TWT).

|  | ABH<br>(year) | DBH<br>(mm) | RW<br>(mm) | WD<br>(kg.m<sup>-3</sup>) | TRW<br>(μm) | TTW<br>(μm) | TWT<br>(μm) |
| --- | --- | --- | --- | --- | --- | --- | --- |
| $\sigma_F^2$ | 7.44 x<br>10<sup>-2</sup> | 3.03 x<br>10<sup>-2</sup> | 5.13 x<br>10<sup>-3</sup> | 2.77 x<br>10<sup>-3</sup> | 5.99 x<br>10<sup>-4</sup> | 4.73 x<br>10<sup>-4</sup> | 2.47 x<br>10<sup>-3</sup> |
| $\sigma_B^2 + \sigma^2$ | 1.65 | 2.15 x<br>10<sup>-1</sup> | 5.63 x<br>10<sup>-2</sup> | 1.95 x<br>10<sup>-2</sup> | 6.04 x<br>10<sup>-3</sup> | 4.91 x<br>10<sup>-3</sup> | 1.88 x<br>10<sup>-2</sup> |
| $h^2$ (%) | 17.31 | 49.32 | 33.47 | 49.80 | 36.11 | 35.20 | 46.52 |

### Accounting for early growth rate ruled out environmental causes from the analysis of ontogeny

By definition, ontogenic tempo is given by the ABH. It had a relatively higher environmental variance, and therefore a relatively lower heritability of 17.31% than other traits (Table 1). The lower heritability was not due to a lower genetic variance, in fact quite the opposite. ABH also depended on the geographical origin of the genetic cluster, as the genetic cluster effect was significant (χ^2^ = 102.8, df = 1, p < 10^−5^ in model (1)). As mentioned earlier, ABH was latitudinally structured, half-sib families of southern origin having low ABH reflecting higher early growth rates.

Despite evidence supporting the genetic control of ABH, the genetic signals in the model (2) seems to have been absorbed by the genetic cluster and the family effects, and let the feature “ABH” only reflect environmental factors out of a lesser correlation with the response variable: when correcting for multiple testing, in model (2), the effects of ABH and of the genetic clusters were significant for more than 70% of the phenotypes (47 out of 66 for both), whereas their interaction was significant for less than 2% of the phenotypes (1 out of 66; Table S2). In other words, ABH explained a large part of the phenotypic variation, while being orthogonal to the family or genetic cluster effect - the variable “ABH” was then likely only capturing the environmental part of ABH. In this particular statistical context, we will assume that ABH reflects environmental factors – else than the block effect already accounted for – impacting early growth and orthogonal to genetic factors (e.g., microenvironmental heterogeneity of the field, supposedly orthogonal to genetic factors because of the random block design). Under this assumption, it is possible to disentangle the effect of genetic factors on the phenotype of a “mature” tree from environmental factors giving advantages in early stages of life (i.e., acclimation). We will come back to this interpretation in the Discussion.

### Early favorable environment mainly impacted macro-phenotypes

The effect of acclimation, approximated by ABH, varied mainly with the cambial age, following a consistent pattern for every trait (but RW): acclimation mostly impacted early (3^rd^ and 4^th^) or later (9^th^ and higher) rings, and least impacted intermediate rings (5^th^ to 8^th^). More precisely, analysis of variance of model (2) showed that, for early rings, ABH significantly impacted every trait but the 4^th^ ring WD (χ^2^ = 7.60, df = 1, p = 5.84 × 10^−3^). For intermediate rings, ABH significantly impacted only 6 out of 20 traits, namely the 5^th^ ring DBH, the 6^th^ ring TTW, the 7^th^ ring TTW, and the 8^th^ ring DBH, WD, and TWT (χ^2^ > 13.9, df = 1, p < 1.88 × 10^−4^). For later rings, ABH significantly impacted every trait but the 9^th^ ring TTW (χ^2^ = 3.30, df = 1, p = 6.92 × 10^−2^), the 10^th^ ring TTW (χ^2^ = 4.70, df = 1, p = 3.02 × 10^−2^), the 11^th^ ring TWT (χ^2^ = 5.65, df = 1, p = 1.75 × 10^−2^) and the 12^th^ ring TWT (χ^2^ = 9.98 × 10^−2^, df = 1, p = 0.752). Only RW followed another pattern, for which ABH impacted mostly intermediate and later rings (5^th^ ring and beyond), as the significance of ABH followed a concave curve starting with an increase of significance from the 5^th^ ring (χ^2^ = 40.70, df = 1, p < 10^−5^) up until the 9^th^ ring (χ^2^ = 1199, df = 1, p < 10^−5^), then a decrease until the 13^th^ ring (χ^2^ = 54.50, df = 1, p < 10^−5^). This can also be seen in Fig. 5a of Lundqvist et al. (2018), where similar age curves, shifting sidewise with ABH, have their steepest phase during the intermediate cambial ages (see also Supplemental Material, method A).

On later rings, trees with a low ABH, i.e., fast growers, ended up with a markedly larger DBH, lower WD and TWT, and slightly higher TRW and TTW. Fast growers had increasingly higher DBH over time, as the effect of ABH decreased from the 3^rd^ ring to the 13^th^ ring (from 5.36 × 10^−3^ to −2.87 × 10^−2^). On the opposite, fast growers had increasingly lower WD, as the effect of ABH increased from the 3^rd^ ring to the 13^th^ ring (from 4.93 × 10^−3^ to 3.1 × 10^−2^). Fast growers also had significantly larger 9^th^ rings, as the effect of ABH on RW followed a convex curve matching its concave significance, reaching its largest negative value at the 9^th^ ring (−5.82 × 10^−1^). At all cambial ages, the effect of ABH was smaller for tracheid properties (TRW, TTW, TWT) than for macro-phenotypes (DBH, WD) (Fig. S2). These results show that environmental factors favoring early growth had an impact on all later stage phenotypes (except for RW), albeit stronger on macro-phenotypes (DBH, WD) than on micro-phenotypes (TWT, TRW, TTW).

### Macro-phenotypes were also the ones showing signature of past divergent selection

In addition to all phenotypes being, as shown above, significantly associated with geographical location, the significance of this association increased over cambial ages (Student’s t > 4.36, p < 0.002), apart from TRW (Student’s t = −0.764, p = 0.464). The significance of the genetic cluster in model (2) on the 13^th^ ring was large for DBH and RW (χ^2^ > 131.26, df = 5, p < 10^−5^), intermediate for WD and TWT (χ^2^ > 79.51, df = 5, p < 10^−5^), and low for TRW and TTW (χ^2^ < 19.39, df = 5, p < 0.002). Presumably, the significance of TWT might be due to its high correlation with the significant WD (ρ = 0.88). The variation of growth-related macro-phenotypes, namely DBH and RW, therefore reflected the population structure more pronouncedly than did tracheid dimensions or wood density (Fig. 2A).

**Figure 2.**
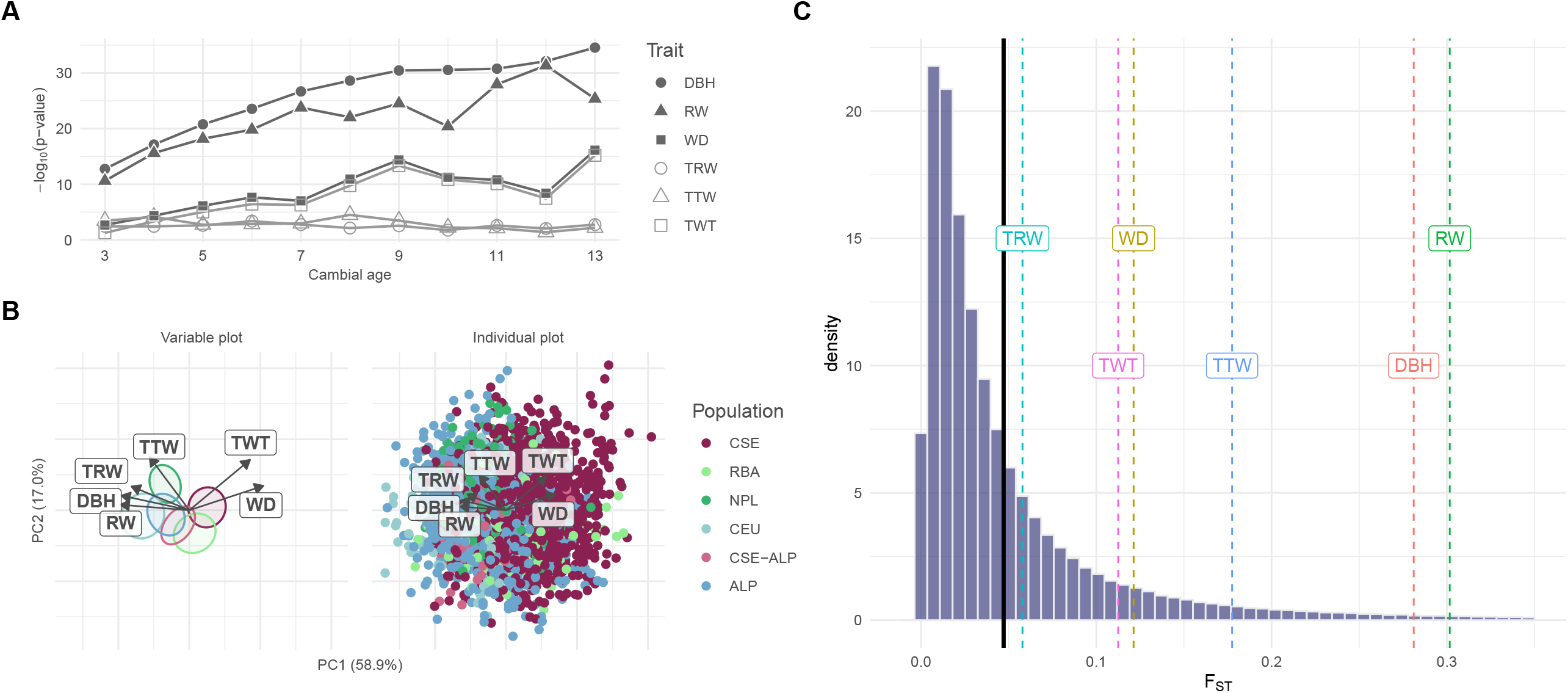
Genetic control of wood properties in Norway spruce. **(A)** ANOVA of genetic clusters in model (2) as a function of cambial age (from 3 to 13), and of traits: DBH (plain circle), RW (plain triangle), WD (plain square), TRW (circle), TTW (triangle), and TWT (square). **(B)** Principal Component Analysis of the residuals of the phenotypes according to model (2): Diameter at Breast Height (DBH), Ring Width (RW), Wood Density (WD), Tracheid Radial Width (TRW), Tracheid Tangential Width (TTW), and Tracheid Wall Thickness (TWT). Left panel only shows variable axes and 10% confidence ellipses (zoomed x2.5), where clusters were ordered in decreasing latitude from right to left. On the right panel, each data (dot) is colored according to its genetic cluster (ordered by decreasing latitude in the legend): Central and South Sweden (CSE, dark red), Russia-Baltics (RBA, light green), Northern Poland (NPL, green), Central Europe (CEU, light blue), Alps (ALP, blue), and hybrids between CSE and ALP (CSE-ALP, pink). PC1 and PC2 explained 57.5% and 18.2% of the variance, respectively. **(C)** Distribution of pairwise F_ST_ across the genome, averaged among pairs of populations. The vertical solid black line is the average F_ST_ over the genome (0.0488), and vertical dotted lines are the Q_ST_ estimated for different traits (averaged across rings): Diameter at Breast Height (DBH, red, Q_ST_ = 0.281), Ring Width (RW, green, Q_ST_ = 0.302), Wood Density (WD, brown, Q_ST_ = 0.121), Tracheid Radial Width (TRW, turquoise, Q_ST_ = 0.0579), Tracheid Tangential Width (TTW, blue, Q_ST_ = 0.177), Tracheid Wall Thickness (TWT, purple, Q_ST_ = 0.112), and Age at Breast Height (ABH, pink, Q_ST_ = 0.066).

Population structure was even more visible on phenotypes when correcting for environmental factors, as the PCA on family effects clearly exhibited the population structure (Fig. 2B): genetic clusters were disposed along PC1 (58.9% of variance explained) in decreasing latitudinal order, except for CEU and ALP being reversed. PC1 was mostly driven by DBH, RW, TRW and WD, while PC2 (17.0% of variance explained) was mostly driven by TTW and TWT. That the traits are structured by both family and genetic cluster can also be seen in the fact that intra-family (or intra-population) variance was significantly increased when the family (or population) structure was shuffled (see Supplement, method B). Not only did phenotypic variation reflect the underlying population structure, but it was also quantitatively associated with latitude.

The quantitative association between environmental gradients and phenotypic variation was partly due to selection (Fig. 2C), at least for macro-phenotypes. Selection as assessed by Q_ST_ from model (3) was only significant for DBH (p = 0.0115), RW (p = 0.0086), and marginally so for TTW (p = 0.0437). Despite the Q_ST_ of TTW being marginally significant, what is striking is that TRW, which is a component of DBH and RW, is not (p = 0.2680). Although interpreting the difference between Q_ST_ and F_ST_ can be fraught with difficulties, one could note that whatever the distribution of F_ST_, selection, if any, would primarily act upon growth-related macroscopic traits. This strong association between population structure and signatures of selection made impossible the inference of genetic variants underlying phenotypic variation since correction for population structure in GWAS made all associations non-significant (see Supplement, method C; Fig. S3-S4). Overall, phenotypic divergence in growth-related macroscopic traits (DBH and RW) is partly due to divergent selection on top of neutral divergence.

## Discussion

The Swedish breeding program is a mixture of local and recently introduced trees from other European countries. As such, although incomplete, the progeny trials reflect the population structure due to glaciation events and constitute an important source of information for evolutionary biology (Chen et al., 2019; Milesi et al., 2019; Li et al., 2022). The breeding conditions constituted a unique opportunity to accurately correct phenotypes for environmental factors, by ruling out recruitment selection and considerably lowering interspecific competition. Furthermore, by correcting for population structure and using common garden experiments, we can rule out neutral and plastic processes so that the observed phenotypic differences can be interpreted as being the outcome of adaptation (Savolainen et al., 2013; de Villemereuil et al., 2020).

In our common garden experiments, growth exhibited a high level of population differentiation, as expected (Savolainen et al., 2007; Kremer et al., 2012). Population differentiation followed a latitudinal pattern, as previously reported (Rossi et al., 2015; Montwé et al., 2016; Milesi et al., 2019). The growth rate was generally differentiated among individual trees at early stages, supporting the fact that young ages are the most dynamic phases (Lundqvist et al., 2018). Differences in apparent growth rate were mainly due to a longer growth span (see discussion in Milesi et al., 2019), and despite its low heritability, its high genetic and environmental variances suggest that it could be related to fitness (Houle, 1992; Caballero, 2020, p.235). This is also supported by the fact that the effect of ABH (here only capturing the micro-environmental effect) was strongest on phenotypes on which signature of selection was also the strongest. Early advantages, possibly acquired by chance, seem to be re-invested by default on resultants – traits giving competitive advantages.

### The quantitative association between phenotypes and population structure predates the Last Glacial Maximum

Phenotypic variation had a clear latitudinal component at both genetic and phenotypic levels, as previously reported in Norway spruce (e.g., Milesi et al., 2019) and in other species (e.g., Sang et al., 2019; Csillery et al., 2020). Since local adaptation matches the environment (Savolainen et al., 2013) and post-glacial recolonization routes followed environmental gradients (Gaggiotti et al., 2009; Li et al., 2022), population structure of Norway spruce is, as a result, strongly confounded with local adaptation. Population structure is thus a good predictor of selection in Norway spruce, as previously reported (Collignon et al. 2002; Savolainen et al., 2007; Kremer et al. 2010). However, local adaptation is not necessarily limited to recent population history. Some of the adaptations displayed by Norway spruce were associated to ancestral environments, and some to current environments. If we consider that adaptation can be dated by the population structure it is associated with, knowing that the population structure of Norway spruce largely predates the Last Glacial Maximum (10 to 15 million years ago; Chen et al., 2019; Li et al., 2022), the observed phenotypic divergence is partly due to adaptations also predating the Last Glacial Maximum. We posit that the quantitative association between phenotype, latitude, and population structure would not be observed if there were no local adaptation to ancestral environments. The last post-glacial expansion of Norway spruce was thus mainly shaped by adaptations *formerly* acquired – not *newly* acquired. In other words, the phenotypic variation within the refugia explains in part the latitudinal variation observed nowadays at large geographical scale.

Because of the association between phenotypic variation and demographic processes, assessing the genetic control underlying adaptation of Norway spruce is challenging: not correcting for population structure in GWASs, will lead to a high false positive rate (Barton et al., 2019; Lawson et al., 2019), and correcting for population structure, may remove true association, i.e., lead to a high false negative rate. A comprehensive assessment of the genetic variants underlying phenotypic variation is then not possible when there is such a strong latitudinal component. Increasing the number of samples or the number of SNPs remains inconclusive (Chen et al., 2021). A possible workaround suggested by Milesi et al. (2019) is to consider phenotypes that are less latitudinally associated, such as the longitudinally associated bud burst, and correct other phenotypes by the association with population structure thereby inferred. Alternatively, we can consider several species jointly as suggested by Alberto et al. (2013).

### Divergent selection mostly affected resultants, not components

The strongest genetic divergence was observed for growth-related traits, and secondly for wood density, suggesting that most of the adaptation is associated with radial or upright growth. Divergent selection acting at early life stages most likely led to the accumulation of competitive advantages, causing then an observable ontogenetic shift of the effect of population structure. The average value of F_ST_ was slightly lower than previously reported (0.0488 in our study, and ∼0.116-0.145 in Table 1 of Chen et al., 2019), and the interpretation of Q_ST_ *vs* F_ST_ remains difficult (e.g., Whitlock, 2008; Edelaar et al., 2011). However, overall, correcting the estimates of F_ST_ would not change the conclusion of our study: if there is selection, it would affect growth-related macroscopic traits more strongly than wood density or its microscopic components. Consistently, the PCA on family effects showed that the latitudinal component – and so, population structure – was mostly explained by stem diameter, ring width and wood density. Neutral divergence could not be excluded for early growth rate (ABH) in spite of its clear relation with growth. Either this trait is neutral, which is highly unlikely as early-stage selection is known to be strong (e.g., Saleh et al., 2022), or, more likely, selection on ABH is invariant among genetic clusters. In the latter case, selection on early growth rate would be strong regardless of the heterogeneity of the locations, where variability would not come from divergence but from, for instance, trade-offs with other life-history traits (e.g., wood density).

The difference between Q_ST_ and F_ST_ is due to the covariance between effect sizes, and the covariance between allelic frequencies (reviewed in Le Corre & Kremer, 2012). Altogether, the resulting pattern of our analyses is that adaptation is more tightly associated to traits that are resultants rather than to its components, in agreement with Caballero (2020, p. 253). Indeed, components can have random variations that are invisible to selection – therefore neutral, as long as the components contribute to a resultant that is fit. In other words, components have a higher degree of freedom, and therefore are noisier. Since genetic polymorphism depicted by SNPs are partly linked to selection, genomic selection on resultant – such as stem diameter – should be much more efficient than on components – such as tracheid traits. An alternative explanation is simply that components were not associated with divergent selection predating the Last Glacial Maximum, and might be under recent selection – not detectable with our approach.

## Conclusion

As expected, phenotypic variation reflected the population structure of Norway spruce. The fact that the phenotypes were latitudinally ordered suggests that the origin of population was strongly confounded with adaptation. Teasing apart the demographic history and the adaptation history is challenging, more so that it has been demonstrated that population history had a strong impact on phenotypic divergence in Norway spruce (Lagercrantz & Ryman, 1990; Milesi et al., 2019). Divergence of phenotypic traits might in part be the results of past adaptations, which can be dated by the population structure it is associated with. In accordance with previous studies (Savolainen et al., 2013; Milesi et al., 2019; Martinez-Sancho et al., 2021), our results support the hypothesis that the current distribution of Norway spruce reflects in part adaptations to ancestral environments, as some signatures of selection were associated with population structure that predated the Last Glacial Maximum. The divergent selection affected more growth-related macro-phenotypes than microphenotypes at tracheid level – presumably invisible to selection.

## Author contribution

RGG, SOL, and BK designed the original study; TG, LO and SOL managed the phenotypic characterization of the samples and organized the data; PM, ML and RGG contributed genomic data; MT and PM analyzed the data; MT, PM, ML, SOL, RGG drafted the manuscript. All authors read and approved the final version of the manuscript.

## Data Availability

The exome capture raw reads and the RNA-seq data have been deposited in NCBI’s sequence read archive (SRA) under accession number PRJNA511374 (Chen et al., 2019) and PRJNA731384 (Chen et al., 2021). For question on availability of the raw phenotypic data, contact RGG.

## Acknowledgements

The authors acknowledge the Swedish research program Bio4Energy and Skogforsk for access to phenotypic data, and the Swedish Foundation for Strategic Research (SSF), grant number RBP14-0040 for exome capture data. This study was funded by the following sources: Uppsala University, SLU Umeå, and the European Union’s Horizon 2020 B4Est for basic functioning and postdoc grant for MT. The authors would also like to thank Sophie Karrenberg and Zhi-Qiang Chen for their useful feedback. The computations and data handling were enabled by resources provided by the Swedish National Infrastructure for Computing (SNIC 2017-7-296) at UppMax partially funded by the Swedish Research Council through grant agreement no. 2018-05973.

## Notes

### Competing Interest Statement

The authors have declared no competing interest.

### Summary of Updates

The discussion has toned down its speculation about competition, and refocused on what was shown by the data.

