## Supplementary materials for "Divergent selection predating the Last Glacial Maximum mainly acted on macro-phenotypes in Norway spruce"

### Figures

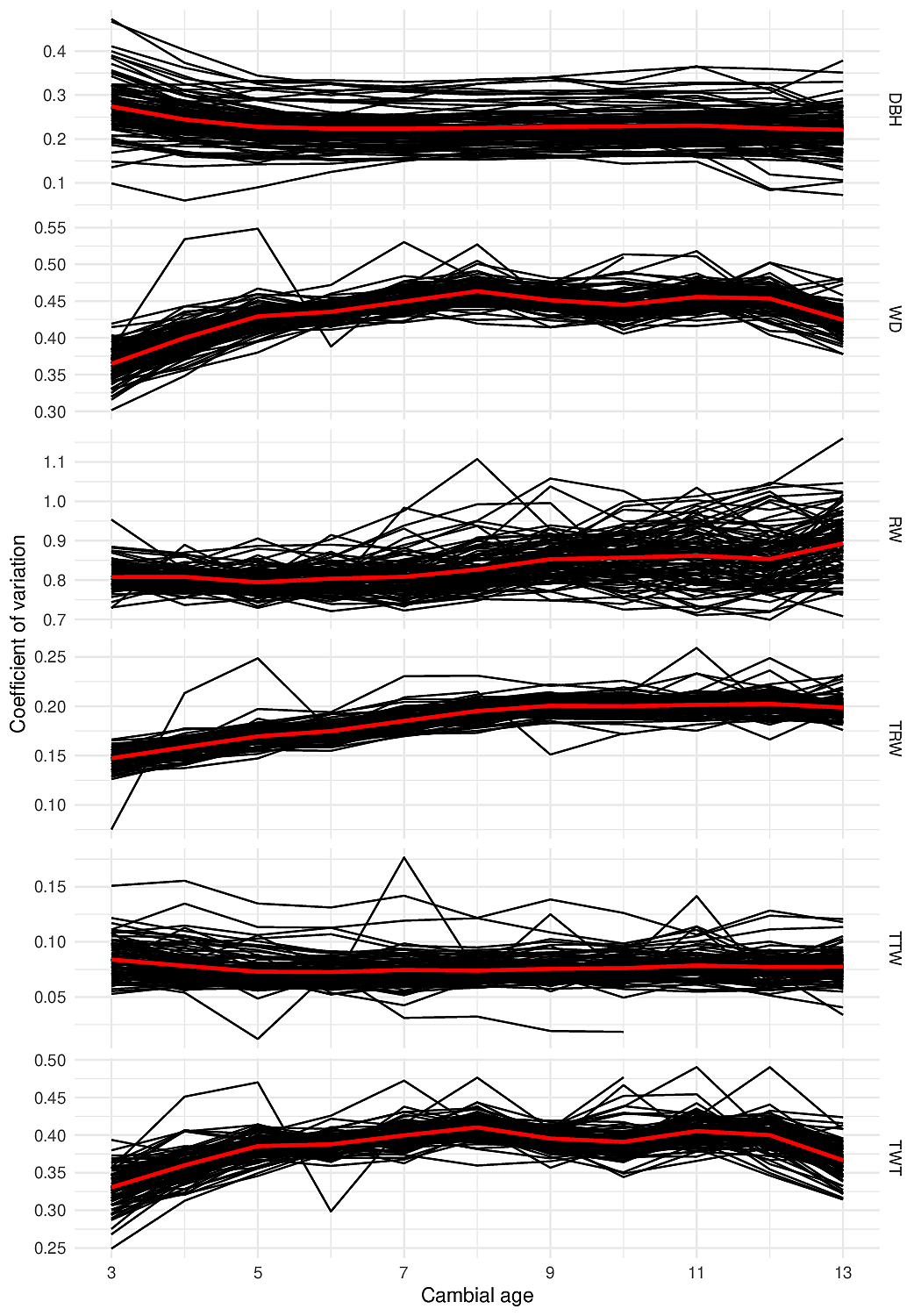

**Figure S1.** Coefficient of variation over cambial age in Höreda. Diameter at Breast Height (DBH), Ring Width (RW), Wood Density (WD), Tracheid Radial Width (TRW), Tracheid Tangential Width (TTW), and Tracheid Wall Thickness (TWT). The black solid curves are the coefficients of variation within one block, and the red solid curve is the average coefficient of variation.

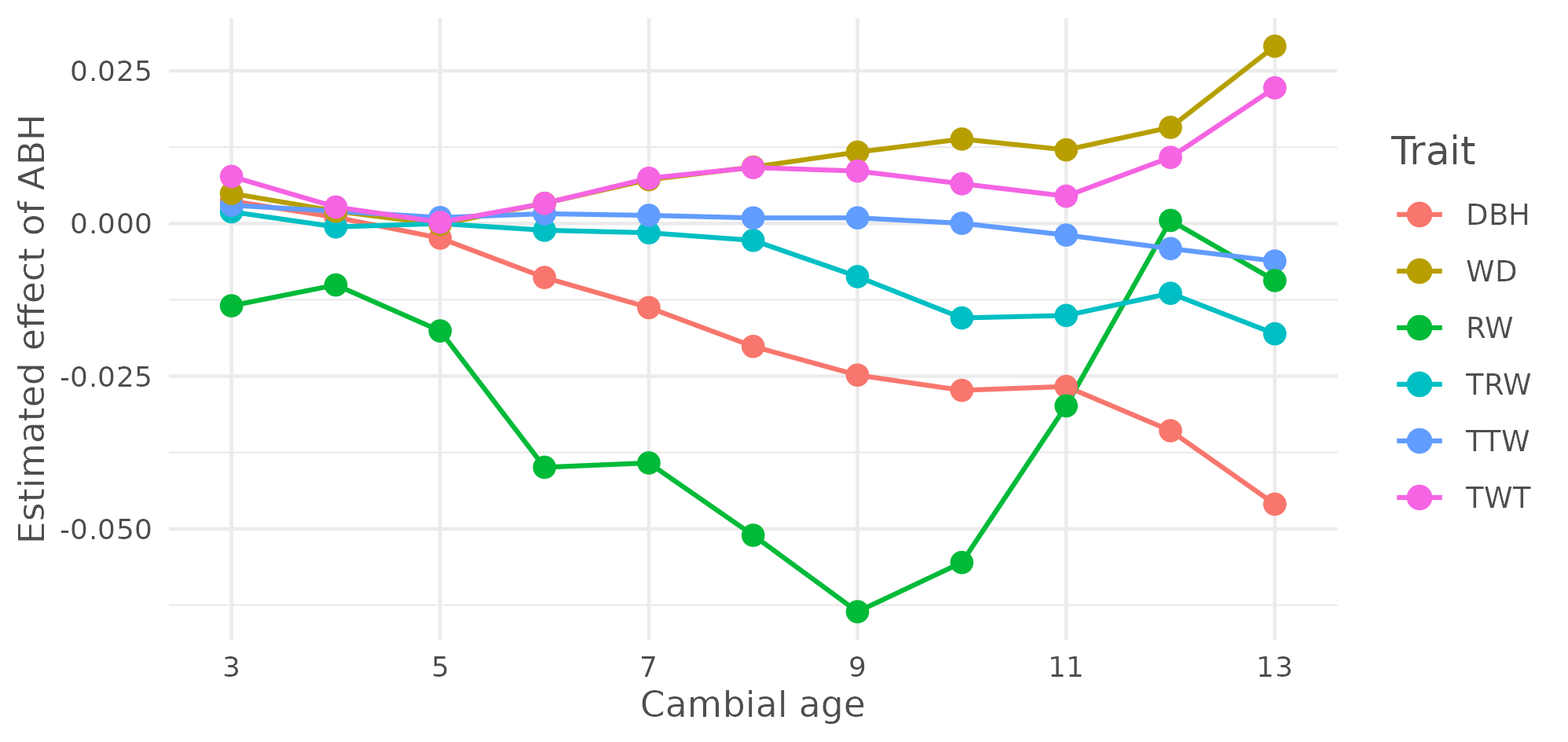
**Figure S2.** Effect of ABH estimated in the model (2) per cambial age (x-axis) and per trait: Diameter at Breast Height (DBH, red), Ring Width (RW, brown), Wood Density (WD, green), Tracheid Radial Width (TRW, turquoise), Tracheid Tangential Width (TTW, blue), and Tracheid Wall Thickness (TWT, pink).

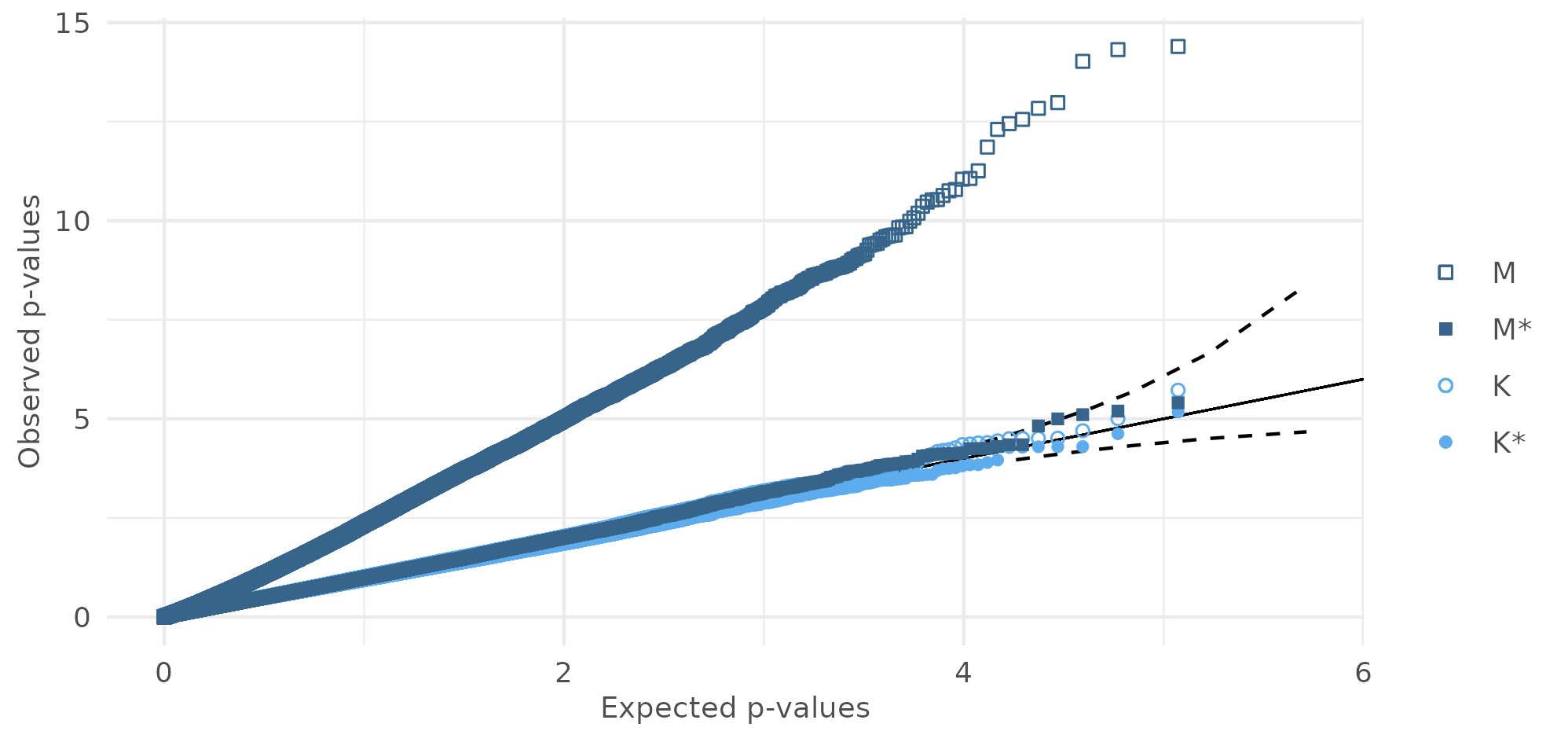
**Figure S3.** Q-Q plot of observed *versus* expected *p-*values (in log scale) on the 5^th^ ring DBH, without any correction (M, in squares), with population structure correction (M*, in plain squares), with kinship correction (K, in circles), and with population structure and kinship corrections (K*, in plain circles). The solid line is a 1-slope line, and the dashed lines are the upper and lower bound of the 95% confidence interval.

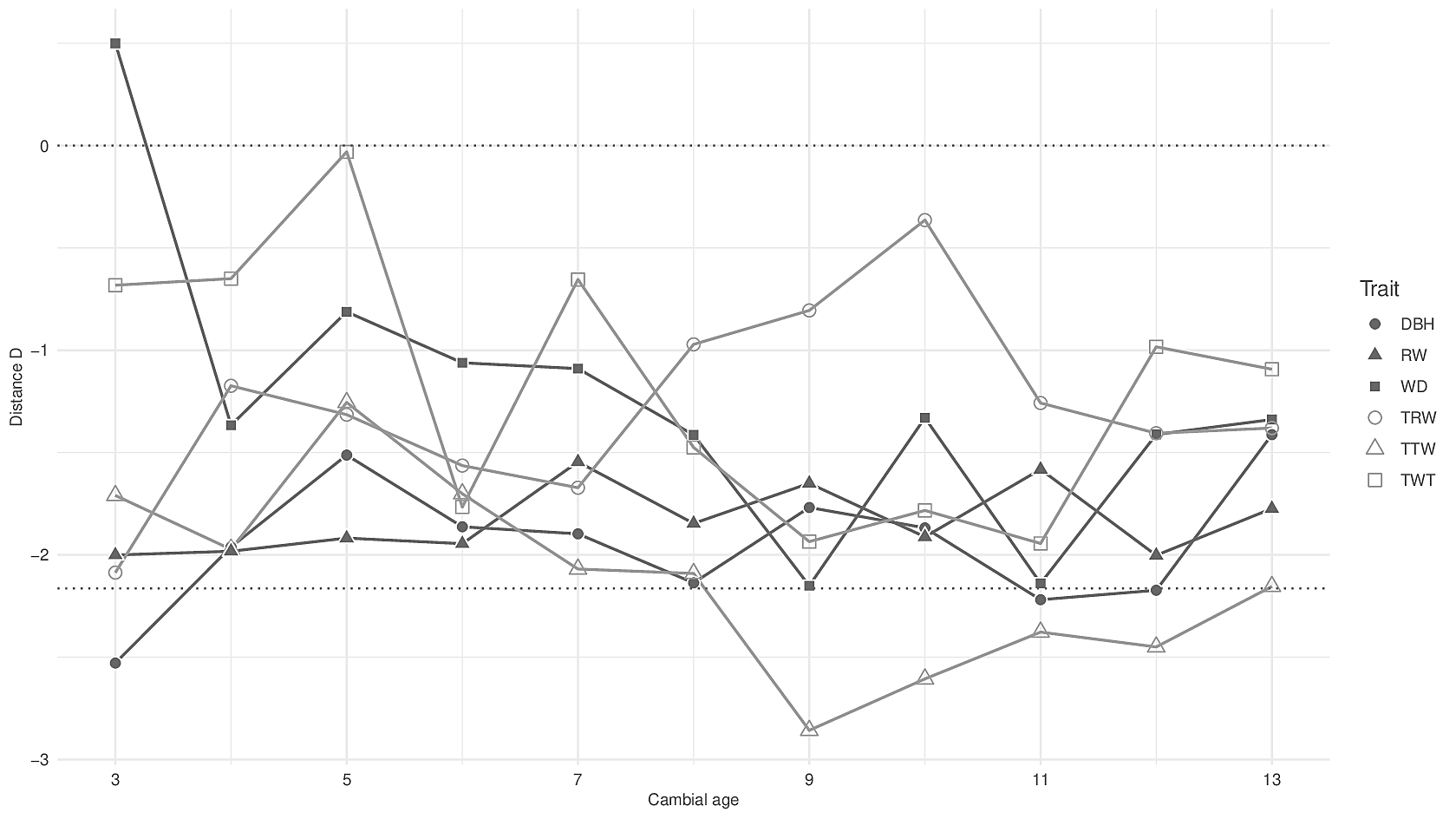
**Figure S4.** Distance *D* on GWAS (K) within the Central and South Sweden (CSE) genetic cluster as a function of cambial age and of traits: DBH (plain circle), RW (plain triangle), WD (plain square), TRW (circle), TTW (triangle), and TWT (square). The dotted horizontal lines indicate the width of the 95% confidence interval.

### Tables

| **Genetic cluster** | **Number of families** |
| --- | --- |
| CSE | 264 (206) |
| RBA | 25 (21) |
| NPL | 32 (27) |
| CEU | 20 (18) |
| ALP | 14 (10) |
| CSE-ALP | 117 (111) |

**Table S1.** Number of half-sib families per genetic cluster, and in parentheses the same number after applying all the filters (summing to 396 families). Genetic clusters: Central and South Sweden (CSE), Russia-Baltics (RBA), Northern Poland (NPL), Central Europe (CEU), Alpines (ALP), and hybrids between CSE and ALP (CSE-ALP).

|  | **df** | **n*** | **DBH** | **RW** | **WD** | **TRW** | **TTW** | **TWT** |
| --- | --- | --- | --- | --- | --- | --- | --- | --- |
| **c** | 1 | 47 | 62.19*** | 341.73*** | 46.14*** | 169.27*** | 17.63*** | 29.24*** |
| **s** | 1 | 52 | 59.78*** | 47.38*** | 48.94*** | 41.16*** | 24.87*** | 43.82*** |
| **g** | 5 | 47 | 132.46*** | 111.74*** | 50.76*** | 17.79** | 19.96** | 45.89*** |
| **c x s** | 1 | 32 | 17.52*** | 41.30*** | 30.91*** | 23.31*** | 1.50^n.s.^ | 39.87*** |
| **c x g** | 5 | 1 | 11.29* | 4.92^n.s.^ | 9.44^n.s.^ | 11.62* | 10.64^n.s.^ | 8.13^n.s.^ |

**Table S2.** Analysis of variance in model (2) of the effects of Age at Breast Height (ABH) (c), site (s), genetic cluster (g), interaction between ABH and site (c x s), and interaction between ABH and genetic cluster (c x g) on the Diameter at Breast Height (DBH), Ring Width (RW), Wood Density (WD), Tracheid Radial Width (TRW), Tracheid Tangential Width (TTW) and Tracheid Wall Thickness (TWT). Degree of freedom (df), number of significant phenotypes out of 66 at a Bonferroni adjusted cut-off of 7.6 x 10^-4^ (n*), and statistics (χ^2^, averaged across rings) are also reported. Significance levels are: p*** < 0.001; p** < 0.01; p* < 0.05; p^n.s.^ > 0.05.

### Methods

##### A) Competition and the S-shaped growth

In our experiment, early competition was a punctual event, and caused a punctual increase in the annual ring width in fast growers. The effect of ABH on ring width can be viewed as the derivative of that of DBH, where the starting point of the competition matches the inflexion point of the S-shaped growth.

##### B) Intra-family and intra-population variance

Family structure was very strong for every phenotype except for TTW, as the intra-family variance was always lower than simulated variance from shuffling the family structure (n = 1000 times). For TTW however, a large portion of simulated variance was lower than the real intra-family variance in early rings (87.9% of the values at the 3rd ring, 15.3% at the 4th ring, and 0.3% at the 5th ring). In contrast, population structure was more time-dependent, as simulated intra-population variance (n=1000 times) was lower than the real value at least 15.3% of the times in early rings (3rd, 4th and 5th), but at most 5% of the times in later stages rings (from 7th ring). Ring width and TTW were exceptions: for ring width, the number of simulated intra-population variance followed the same pattern as Fig. S2 (a decrease down to 7th ring, then increase); and for TTW, at least 7.9% of simulated values were lower than the real variance.

##### C) GWAS

GWAS was performed on the estimated family effect of model (1) and model (1bis) (model (1) with the genetic cluster effect). We fitted each SNP individually, with or without a population structure correction by including the genetic cluster as a covariate (model (1) or model (1bis) respectively). We also used the software GCTA (option --mlma; Yang et al., 2011) that included a polygenic effect. GWASs not accounting for kinship (model 1 and 1bis) were referred to (M) when not corrected for population structure, and (M*) when corrected for population structure. GWASs accounting for kinship (from GCTA) were referred to (K) when not corrected for population structure, and (K*) when corrected for population structure. GWAS (K*) would be the classic GWAS. For significant SNPs, we re-estimated the effect size by using an empirical Bayes approach for adaptive shrinkage with the R package *ashr* (function *ash*; Stephens, 2017). In order to assess the quality of a GWAS through its Q-Q plot, we introduce the distance D as the distance between the highest log p-value (the rightmost value on a Q-Q plot) and the highest upper bound of the 95% confidence interval of a uniform distribution. If the distance D is positive, it means that some SNPs are more significant than expected; if D is between 0 and −2.16 (the width of the confidence interval for n = 118,145), the p-value distribution follows a uniform distribution with a confidence level of 95%; and if D is below −2.16, the SNPs are less significant than expected, suggesting an overcorrection by population structure. Additionally, in order to detect regions with high density of significant SNPs - namely peaks - we used a custom peak detection script with default settings (Tiret & Milesi, 2021), which is freely available at https://github.com/milesilab/peakdetection (see details in Li et al., preprint).

If selection on quantitative traits acts as a shift of allele frequencies of Quantitative Trait Loci (QTL), a genome scan would detect signature of such selection. However, in the GWASs on the 66 traits, population structure was a confounding factor with genetic control for every single trait, as significant SNPs were removed when correcting for population structure and/or for kinship. When correcting for population structure and/or kinship in the GWASs (M*, K, or K*), estimated p-values followed a uniform distribution with a 95% confidence (Fig. S3, S4), as the distance D was negative for at least 64 out of 66 traits. Only the 8^th^ ring wood density had some SNPs more significant than expected (D = 0.0662), but none of the SNPs were significant after re-estimating the effect size with the adaptive shrinkage framework (*p* > 0.201). As a matter of fact, correcting our sample for population structure or kinship flattened, or even overcorrected, the genetic signals on the Manhattan plots, since, when no correction was applied, SNPs were more significant than expected for 41 out of 66 traits. GWASs within the largest genetic cluster CSE were also inconclusive, as the distance D was also always negative (except for the 3^rd^ ring WD), with or without correcting for kinship.

##### C) Admixture

The ADMIXTURE plot (Fig. 1C) used a further filtering since missingness was correlated to population structure: we then filtered for 30% individual missingness, and 10% SNP missingness, resulting in 344 half-sib families and 58 194 SNPs. The number of families per genetic cluster also varied: CSE = 193, RBA = 16, NPL = 24, CEU = 16, CSE-ALP = 10, ALP = 85.

##### D) Correlation between estimated p-values from different GWASs

The p-values estimated by GWAS (K) and GWAS (K*) (including kinship) were strongly correlated (⍴ > 0.9340). As expected, GWASs (M) and (M*) were strongly correlated when population structure was weak, following a reverse pattern of significance of the genetic cluster (Fig. 4B). Correlation of between GWAS (M) and (K) (or K*) converged towards 0.20 at the 13th ring. Finally, correlation between GWAS (M*) and (K) (or K*) was constant over time per trait: ⍴ = 0.908 for DBH, ⍴ = 0.0675 for ring width, ⍴ = 0.293 for wood density, ⍴ = 0.300 for TRW, ⍴ = 0.0688 for TTW, and ⍴ = 0.938 for TWT.
